# Weather-related changes in the dehydration of respiratory droplets on surfaces bolster bacterial endurance

**DOI:** 10.1101/2024.02.07.579296

**Authors:** Abdur Rasheed, Kirti Parmar, Siddhant Jain, Dipshikha Chakravortty, Saptarshi Basu

## Abstract

**Hypothesis:** The study shows for the first time a fivefold difference in the survivability of the bacterium *Pseudomonas Aeruginosa* (PA) in a realistic respiratory fluid droplet on fomites undergoing drying at different environmental conditions. For instance, in 2023, the annual average relative humidity (RH) in London (UK) is 71%, whereas in Delhi (India), it is 45%, showing that disease spread from fomites could have a demographic dependence. Respiratory fluid droplet ejections containing pathogens on inanimate surfaces are crucial in disease spread, especially in nosocomial settings. However, the interplay between evaporation dynamics, internal fluid flow and precipitation and their collective influence on the distribution and survivability of pathogens at different environmental conditions are less known.

**Experiments:** Shadowgraphy imaging is employed to study evaporation, and optical microscopy imaging is used for precipitation dynamics. Micro-particle image velocimetry (MicroPIV) measurements reveal the internal flow dynamics. Confocal imaging of fluorescently labelled PA elucidates the bacterial distribution within the deposits.

**Findings:** The study finds that the evaporation rate is drastically impeded during drying at elevated solutal concentrations, particularly at high RH conditions. MicroPIV shows reduced internal flow under high RH conditions. Evaporation rate influences crystal growth, with delayed efflorescence and extending crystallisation times. PA forms denser peripheral arrangements under high evaporation rates and shows a fivefold increase in survivability under low evaporation rates. These findings highlight the critical impact of environmental conditions on pathogen persistence and disease spread from inanimate surfaces.

**Graphical abstract:** 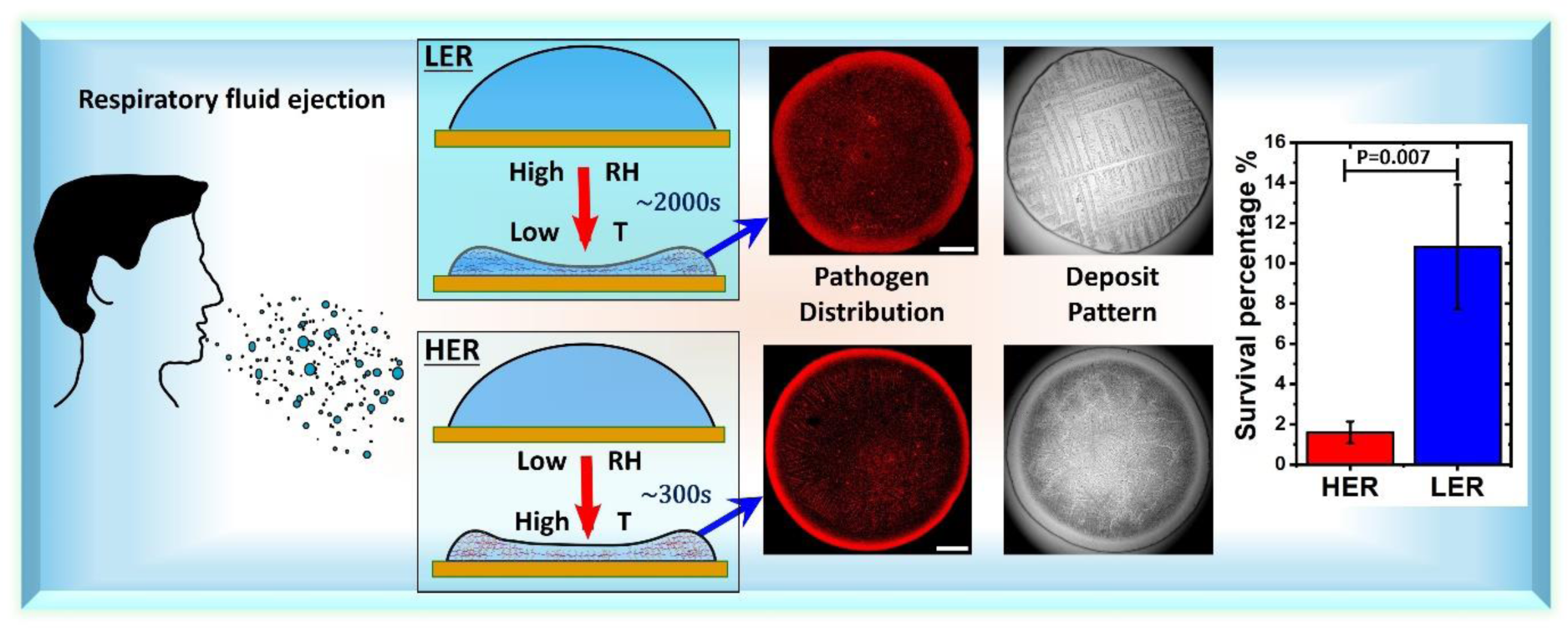

## 1. Introduction

Controlling infectious diseases has been a paramount challenge throughout human history, and the proliferation of diseases has escalated as a significant global concern in the past decade. Ongoing efforts involve implementing various strategies and policies to curb the spread of disease. However, the rise of mutations and antibiotic resistance in pathogens introduces additional hurdles to effective disease control. Furthermore, the survivability and infectivity of pathogens in the environment exhibit species-specific characteristics [1–5].

Bacterial infections, with a staggering global mortality rate of 7.7 million in 2019, emerged as the second leading cause of global deaths, following ischemic heart disease [6]. *Pseudomonas Aeruginosa* (PA) is among the five leading bacteria responsible for global deaths due to bacterial infections [6]. This study focuses on PA, which can infect various organs, including the lungs and bloodstream. PA transmission can be facilitated through respiratory droplets expelled during coughing and sneezing, which contain a complex composition including bacteria, mucin, salt, and other minor compounds[7,8]. These droplets undergo a drying process while suspended in the air or when in contact with surfaces in a sessile mode. As the respiratory droplet undergoes drying, bacteria experience osmotic stress, shear, and desiccation stresses. The dehydration of bacteria during this process is a crucial mechanism leading to their eventual viability.[9] Reducing water content within a bacterial cell can initiate metabolic, structural, and mechanical changes, causing substantial damage to membranes, DNA, and proteins.[10]

Various ambient factors, such as relative humidity, temperature, and ventilation, significantly influence the degree of stress imposed on bacteria [11–15]. Atmospheric relative humidity exhibits variations throughout the day and across different seasons and geographical locations. For instance, in 2023, the annual average of RH in London (UK) is 71%, whereas in Delhi (India) it is 45%. Big droplets settle under gravity in high humidity conditions but remain suspended in the air for an extended duration in dry conditions[16,17]. Distant airborne aerosol dispersal and transmission are mitigated at high relative humidity in classroom settings.[18,19] Multiple studies report on the airborne pathogen survival and transmission [20–28]. On inanimate surfaces, studies [29–32] have explored the survivability of microbes under various environmental conditions, employing culture media such as Luria broth and DMEM for microbial suspension. These studies typically involve large droplet volumes exceeding 10 microliters, atypically larger than those found in sneeze and cough ejections[33–35]. It is important to note that droplet volume changes can influence not only evaporation, precipitation, and subsequent pathogen distribution but also survivability[36]. Studies using droplet volumes of one microliter and less using surrogate biological fluids on different substrate surfaces[37] and different nutrient mediums [38] have been conducted at fixed ambient condition.

A comprehensive understanding of the effect of ambient conditions on pathogen survival and disease transmission from inanimate surfaces using realistic respiratory fluid is still not established. Furthermore, the impact of organic compounds, such as mucin, on the survivability of pathogens in ejected droplets remains less understood. Persons with lower respiratory infections show hypersecretion of mucin up to 9 grams/litre[39]. Studies have shown mucin is hygroscopic in nature and can adsorb and hold moisture[40–42]. Groth et al. showed human respiratory cough droplets containing mucin can grow at a high relative humidity of 90 % by adsorbing water from the ambient. The water adsorption of mucin reduces with relative humidity (RH) [43]. While there are some reports on the effect of the hygroscopic nature of mucin in lung fluid on the evaporation and efflorescence of aerosols[44–47], a comprehensive analysis of the hygroscopic effects of lung fluid on evaporating sessile droplets is not available. Thus, the current study is focused on PA-laden respiratory droplets on inanimate surfaces at low and elevated RH conditions. The respiratory fluid is made in the laboratory using mucin and NaCl. The present study uses porcine gastric mucin-type III, which is shown to have mechanical properties similar to human mucin [8,48]. We hypothesise that the water adsorption at high RH could delay evaporation and reduce bacteria desiccation stress, leading to an increased survival rate. The PA-laden surrogate respiratory fluid (SRF) droplets cast on glass substrate are subjected to two distinct ambient conditions with RH and a temperature varied to maintain low evaporation rate, LER and high evaporation rate, HER conditions. Notably, the droplets exhibit significant variations in evaporation, flow, and precipitation dynamics. The ultimate deposition pattern of pathogens and the crystallisation pattern display substantial differences between the two conditions. Crystal nucleation and growth are much delayed under high RH conditions. The overall physicochemical changes and the stresses imposed on PA under two different conditions varied the bacterial viability by at least fivefold.

## 2. Methods and materials

### 2.1. Ambient conditions and substrate preparation

The experimental setup involves maintaining two different ambient conditions to investigate the effect of ambient conditions under a low evaporation rate and high evaporation rate setup. The RH and temperature T are maintained at 70±3% and 25±1°C for LER and 46±2% and 31±2°C for HER in an inbuilt stability chamber of size 2.5m x 2.5m x 2.5m equipped with temperature and humidity control. Glass coverslips serve as the substrate. They are cleaned by sonicating in water for 3 minutes, followed by sonication with isopropyl alcohol for 3 minutes. The cleaned coverslips are then blow-dried using compressed air. Blow drying with compressed air is compared with compressed nitrogen; noticeable differences are not seen in the wetting properties.

### 2.2. Surrogate respiratory fluid preparation

Surrogate respiratory fluid (SRF) is prepared using mucin (Type III. Sigma Aldrich) and NaCl. Specifically, 9 g/L of mucin (Type III - Sigma Aldrich) and 6 g/L of NaCl are added to distilled water, mechanically stirred at 100 rpm for a minute, and then centrifuged at 2000 rpm for 3 minutes. To create the bacterial-laden SRF, 200 microliters of phosphate buffer saline (PBS) containing PA at a concentration of 10^8 CFU/ml is added to 800 microliters of mucin-NaCl supernatant solution.

### 2.3. Evaporation and precipitation experiments

One-microliter droplets of this bacterial-laden SRF are gently placed onto glass coverslips to study the evaporation, flow, and precipitation dynamics. All experiments were repeated at least three times for each case. Top-view and side-view images are simultaneously captured. Nikon DSLR D7200 camera fitted with a Navitar Zoom lens is used for side view, and a Nikon DSLR D7200 camera mounted on a BX51 Olympus frame using Halogen-based illumination (TH4 200, Olympus) is employed for top-view. The capture rate of 1 fps for HER and 5 fps for LER is used. For the precipitation dynamics study, the instantaneous area precipitated, A_p,_ is evaluated using top-view images by manually tracing the boundaries of the crystallised zones. These sketches are processed in ImageJ to obtain the area of each crystallised zone, which is then summed to determine the precipitated area A_p_ at each instant. The total crystallised area A_t_ only includes regions with crystals. In the case of HER, A_t_ excludes the coffee ring deposits.

### 2.4. Flow experiments

For the Micro-Particle Image Velocimetry (Micro-PIV) analysis, the fluid contained fluorescently labelled polystyrene microparticles with a mean diameter of 0.86 μm. Illumination was provided by a Nd:YAG laser at a wavelength of 532 nm, and particle fluorescence emissions were captured by an Imager Intense Lavison camera. The setup utilised a 10X objective lens, offering a field of view of 600μm x 450 μm and a depth of field of 8 μm. The flow measurements were performed at t/t_f_ ∼ 0.1(where t_f_ is the crystallisation end time) for 5 seconds at a horizontal plane z=50μm from the substrate surface near the droplet’s outer ring. Flow data analysis is conducted using the Davis 8.4 software package.

### 2.5. Confocal Microscopy

The deposits are characterised using Leica TCS SP8 confocal microscope to examine the pathogen distribution. Images are captured at various z planes of 1 μm gap. The superimposed image gives the overall pathogen distribution. The bacteria are tagged with FM-4-64 dye (Thermo Scientific). In the confocal microscope study, the integrated intensity for both cases is approximately the same, indicating similar pathogen concentration and no differential attenuation of fluorescent signal.

### 2.6. Viability study

To study the viability of PA, 30 one-microliter droplets are cast for each repetition. The droplets are allowed to dry and remain exposed to the respective ambient conditions for five hours. Subsequently, the deposits are rehydrated using PBS and collected for plating to assess bacterial viability. Pseudomonas aeruginosa was plated on cetrimide agar, a differential medium that allows identification of Pseudomonas aeruginosa. Survival percentage was calculated by dividing the bacterial colony forming unit (CFU) in the rehydrated deposits by CFU in 30 microlitres of culture without any drying treatment.

## 3. Results and Discussion

### 3.1. Evaporation dynamics

The PA-laden SRF droplets are placed gently on the glass substrate. The SRF droplets exhibited a constant contact radius (CCR) mode of evaporation, as shown in Fig. 1a. It is evident from the side-view images that in high humidity conditions, the droplet takes a longer time to dry. The last column in Fig.1a shows the side-view image during crystallisation. In the case of HER, the crystal growth starts from the droplet periphery. In contrast, the crystal growth starts from the droplet centre in the case of LER. The droplet’s instantaneous volume is calculated using the side view images, assuming a spherical cap shape. The assumption of a spherical cap shape can be justified as the droplet’s characteristic length (the contact radius) is lesser than the capillary length for both cases. The evaporation rate remains nearly constant during the initial stages of evaporation. However, the evaporation rate decreases with time as the vapour pressure above the droplet interface reduces with an increase in solutal concentration. Detecting the volume from side-view images becomes challenging towards the end of evaporation when the droplet becomes too thin and the variation is too small for precise measurement.

**Figure 1.**
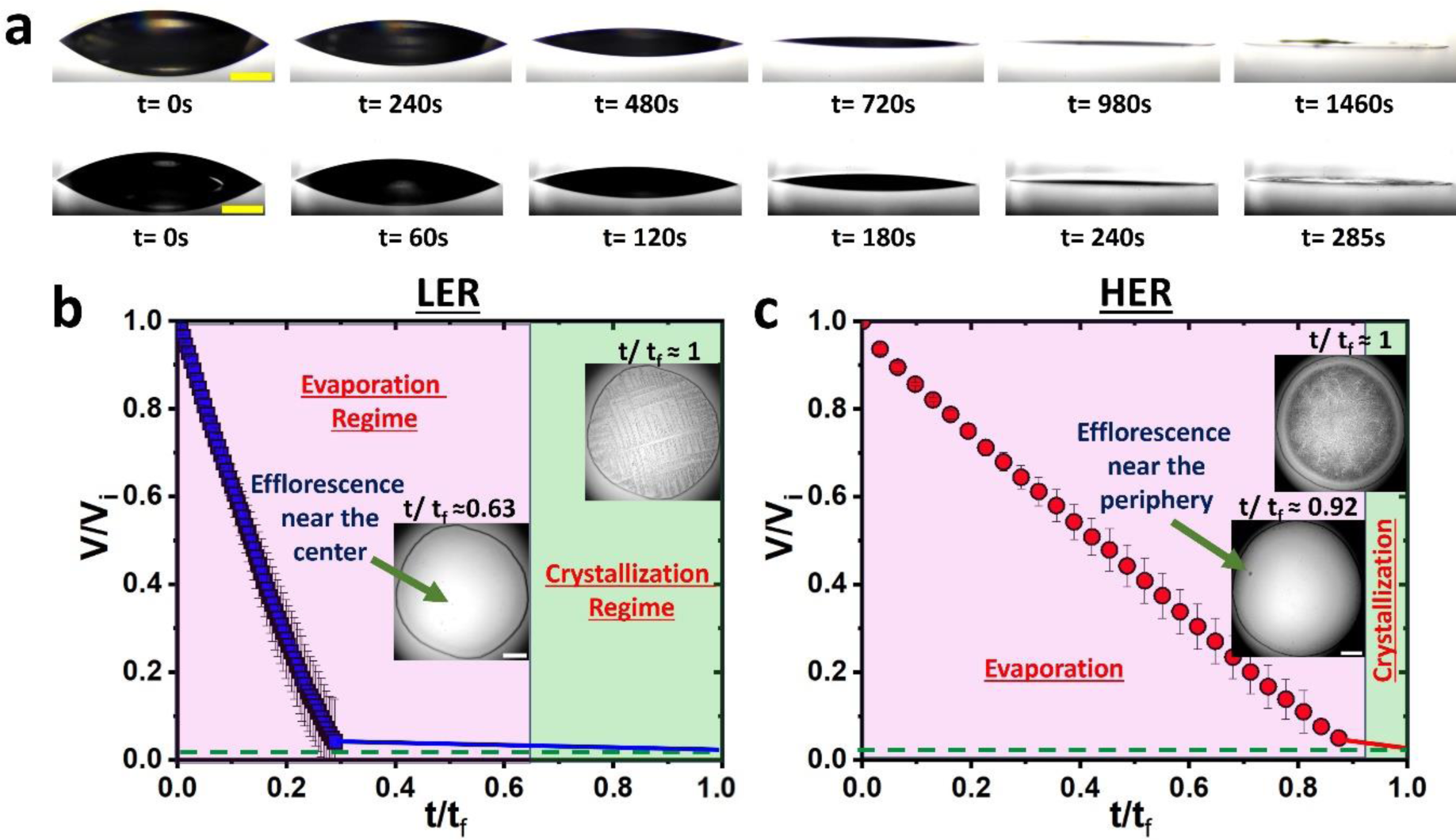
Evaporation dynamics: a) Temporal side-view images of the drying SRF droplet, the top row shows the LER case and the bottom row shows the HER case, scalebar corresponds to 500μm. b) & c) Plot shows the normalised volume, V/V_i_ regression with respect to the time t, non-dimensionalised with t_f_ (time taken till the end of crystallisation) for b) LER case and c) HER case. The plot shows the evaporation and crystallisation regimes with respect to t/t_f,_ and the insets show the top-view images at the onset and the end of crystallisation.

The average evaporation rate of HER case is 3.2e-9 kg/s for the initial volume reduction of 0.965V_i,_ where V_i_ is the initial volume. Under LER conditions, the average evaporation rate is 1.64e-9 kg/s for the initial volume reduction of 0.959V_i_. The lower evaporation rate in the case of LER is attributed to the lesser 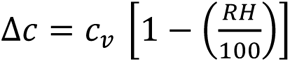, which is the difference in water vapour concentration at the droplet interface compared to the water vapour concentration far away in the surrounding environment[49]. Here, c_v_ is the vapour concentration at the droplet interface, and RH is the percentage of relative humidity. The solutal volume concentration of SRF droplet V_s_/V_i_ is less than 0.015, where V_s_ is the solutal volume. It is difficult to determine if a complete dryout of the droplet has occurred from either side view or top view imaging. The solutal deposits can still hold water and remain hydrated after crystallisation. Thus, the end of crystallisation is considered the end of evaporation and the time taken for crystallisation to end is regarded as the final time t_f_.

Fig1.b & c shows the volume regression with the time non-dimensionalised with respect to t_f_. Concerning the end of crystallisation, LER shows faster volume regression than HER at initial times. The evaporation rate strongly influences the time for the efflorescence of crystals and the crystal growth rate. NaCl typically starts crystal nucleation at a supersaturation level of 2.31 with a saturation concentration of 0.036kg in 0.1kg of water. In the case of HER, crystallisation begins around time fraction t/t_f_ ≈ 0.92, where t=0 s when the droplet is placed on the substrate. For HER, the time for the efflorescence of crystals or crystallisation initiation (t_c_) is 282±9 s, and t_f_ is 316±11 s. Under low evaporation conditions, crystal efflorescence initiates at t/t_f_ = 0.63, with t_c_=1234±97 s and t_f_=1939±106 s. The efflorescence of crystals began in less than 10 s after the volume had reduced to 0.965 V_i_ in the case of HER, whereas it took more than 600 s for the efflorescence of crystals after the volume had reduced to 0.959 V_i_ in the case of LER. It took around 30 s for crystallisation to complete in the case of HER. In contrast, LER crystals took approximately 700 s to grow over the entire wetting area. Such a long delay in efflorescence and crystal growth at LER could be due to the hygroscopic nature [50,51] of mucin.

Figure 2 illustrates the mechanisms behind the observed evaporation and efflorescence phenomena. The proposed mechanism of water sorption and desorption is shown in Figure 2a. Literature shows mucin’s water sorption increases with RH, while desorption reduces with RH[42]. Thus, the water sorption mass flux is lower at HER than at LER. Conversely, the evaporative mass flux is higher at HER than at LER, as shown in figure 2b. The evaporative mass flux is notably higher near the contact line, especially at acute contact angles, as shown in figure 2b. Yet, thick mucin deposits around the droplet edge could lead to a greater water sorption influx at the periphery.

**Figure 2.**
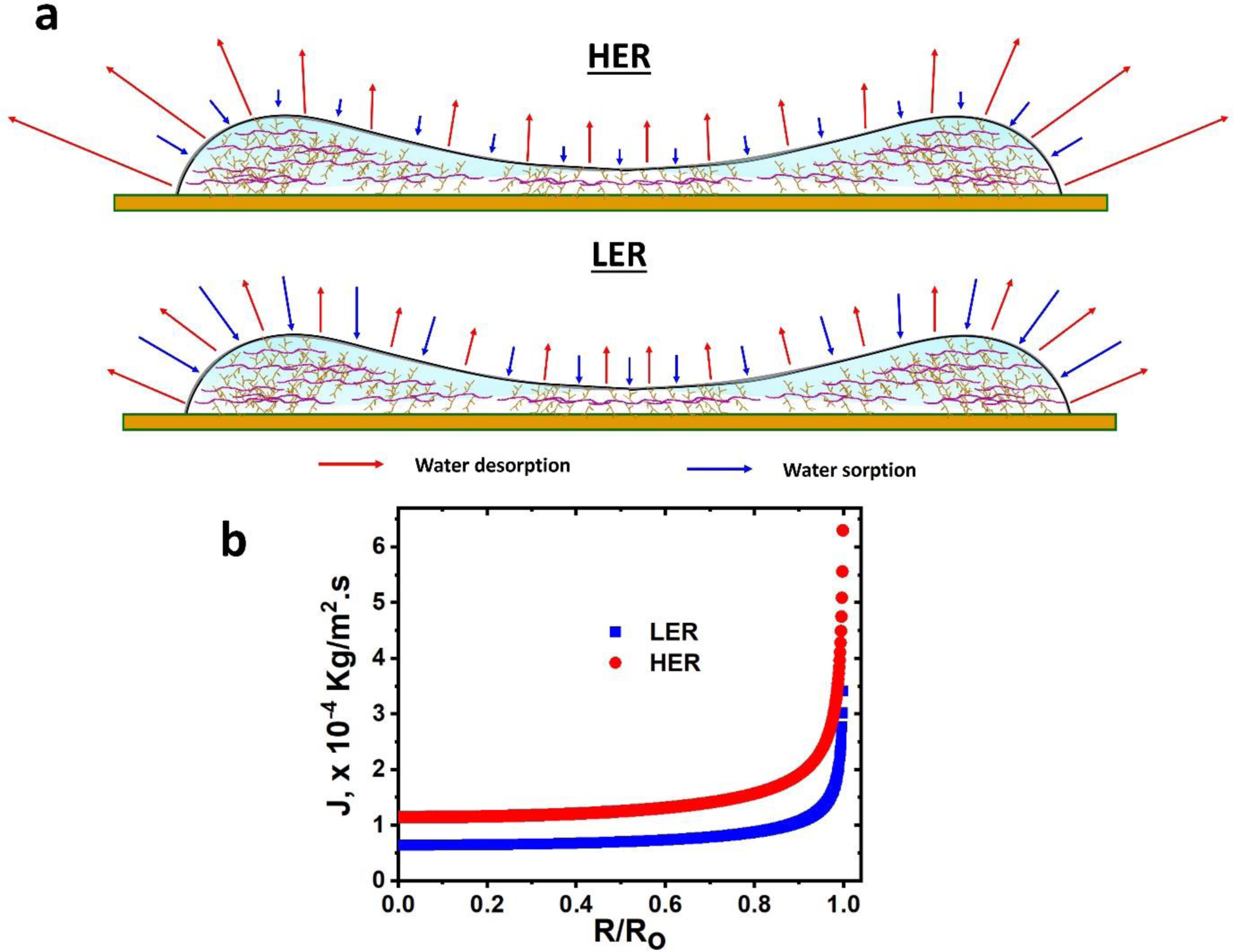
Schematic shows water sorption and desorption fluxes (representative) on mucin deposits at LER and HER conditions leading to crystal nucleation near the periphery in HER and at the central region in LER b) Plot shows the evaporative mass flux variation in the radial direction.

Consequently, the net mass influx and outflux rates can vary across different parts of the droplet. As depicted in Figure 2a, for efflorescence to begin at the periphery under HER conditions, the mass outflux must surpass the mass influx significantly more near the contact line than at other locations within the droplet. At LER conditions, elevated water influx due to adsorption should occur at the periphery compared to the inner regions. Consequently, the central regions might experience a net mass outflux surpassing other areas, leading to crystal efflorescence in the central regions. The evaporative mass flux in a pinned droplet, shown in figure 2b. is calculated using the expression, 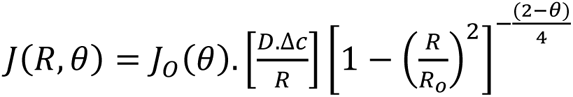,[49] Where, 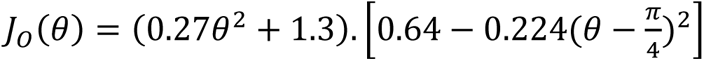,[49]

R is the local radius and R_o_ is the droplet contact radius, D is the water vapour diffusion coefficient in the air in m^2^/s, 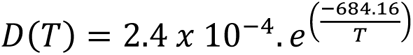[52], and 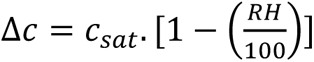, the saturation vapour concentration, c_sat_ =P_sat_/RT, the saturation vapour pressure 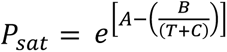, where A, B and C are empirical Antoine constants, A=23.1946, B=3813.98 and C=-46.29 for T in Kelvin.[53] The evaporative mass flux in the case of HER is around twice higher than LER, as shown in Figure 2b.

The volume regression V/V_i_ with time, non-dimensionalised to the characteristic time for diffusive evaporation, τ, is shown in supplementary figure S1a. Here, 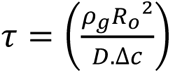 [54,55], where ρ_g_ is the water vapour density. The volume regression of both cases is usually similar, except near the end, where V/V_i_ of the LER case is slightly higher, which could be attributed to water sorption at high RH. The t/τ for efflorescence and crystallisation end is much higher in the case of LER. The volume regression trend in the case of LER is similar to the mass regression curve observed for hygroscopic aqueous droplets[56]. Rigorous investigation using Quartz crystal microbalance with dissipation (QCM-D) technique is required to understand the water sorption and desorption kinetics. Supplementary Figure. S1b depicts variations in t_c_, t_f_, and the contact angle θ. The contact angle experiences an increase of around 1 degree, shifting from 34.7±1.8 to 35.9±1.6 when ambient is changed from HER to LER conditions. According to Young’s equation, 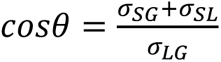, where σ_SG,_ σ_SL_ and σ_LG_ are the relative surface energies of the liquid, substrate, and gas. The variation in surface energies of the SRF fluid and the substrate at different ambient conditions could be responsible for the difference in contact angle. The droplet contact radius remains unchanged till the efflorescence, as shown in supplementary figure S1c. Supplementary figure S1d shows that the contact angle continually reduces as the droplet evaporates. The reduction rate in contact angle is higher for HER than LER due to the high evaporation rate. The evaporation rate influences the desiccation stress on bacteria by affecting the dehydration rate.

### 3.2. Desiccation stress calculations

The degree of desiccation stress on bacterial cells is directly proportional to the rate of dehydration. The rate of dehydration in cells can be given by the rate of change of its water potential (ΔΨ/Δt)[57]. Water potential is calculated as Ψ =P-Π + m_w_gh,[58] where P represents the hydrostatic pressure beyond atmospheric pressure, and Π is the osmotic pressure. The gravitational component becomes relevant mainly at elevated altitude distribution of cells. Thus, the water potential for bacterial cells in tiny droplets simplifies to Ψ =P-Π. The osmotic pressure can be described as Π = Π_s_ - Ψ_m_. where Π_s_ indicates the osmotic pressure from solutes and Ψ_m_ is the matric water potential arising from water’s interaction with surfaces and interfaces. Matric potential becomes significant at low water content (towards evaporation’s end). In this context, water potential is assessed based on osmotic pressure changes within the SRF droplet.

Assuming bacterial cells reach an equilibrium with the SRF during evaporation, their water potential can be considered the same as that of the SRF droplet. The osmotic pressure due to solutes is given by Vant’s Hoff relation Π_s_ =RT Σ_i_ c_i_, with R as the universal gas constant, T is the temperature in Kelvin (299K for LER and 304K for HER), and c_i_ is the concentration of solute species. As evaporation progresses, solute concentration and, consequently, osmotic pressure increases. The variation in water potential through drying is depicted in Figure 3, showing an exponential decrease in water potential as the solute concentration rises. For LER, the reduction in water potential slows at higher solute concentrations due to mucin’s water absorption. Thus, the average dehydration rate is reduced to 0.00893 MPa/s for LER but 0.05301 MPa/s for HER. This analysis provides an approximate estimate of dehydration rates based on solute concentration changes. More detailed calculations involving matric water potential could offer deeper insights into the dehydration rate estimation of bacterial cells. The desiccation stress on bacteria could also vary spatially in the droplet deposition. The droplet deposition is strongly affected by the internal flow dynamics.

**Figure 3.**
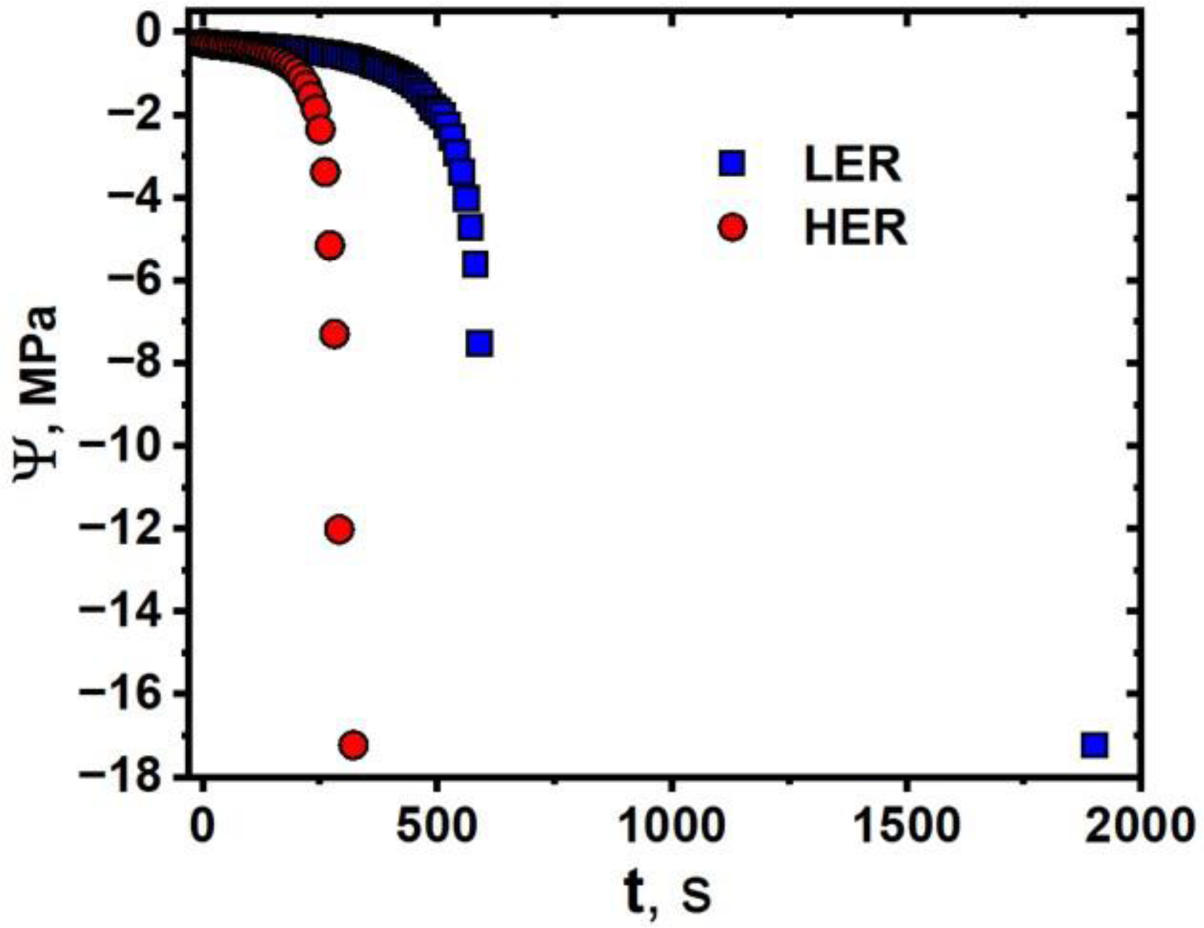
The plot shows variation in water potential in SRF droplet with time for HER and LER cases.

### 3.3. Flow Dynamics

Water loss from the droplet interface induces flow towards the contact line, called capillary flow [59]. In addition, surface tension-driven flows can also occur within the droplet. Evaporative cooling during drying could create surface tension differences due to temperature variations, resulting in thermal Marangoni flow. Solutal Marangoni flow occurs when surface tension is varied by concentration change at the interface. Solutal Marangoni flow is usually found in droplets of miscible binary fluids of different volatilities and salt and surfactant-containing solutions [60–62]. Since the SRF droplet contains dissolved salt, there could be a presence of both capillary and solutal Marangoni flow. Velocity vectors from Micopiv measurements on the SRF droplet at HER and LER conditions are shown in Fig 2. a&b. The planar velocity measurements are made at approximately 50 μm from the substrate surface near the droplet periphery, as shown in figure 4a. The low velocity magnitudes indicate predominantly capillary flows, while the Marangoni flows seem to be suppressed.

**Figure 4.**
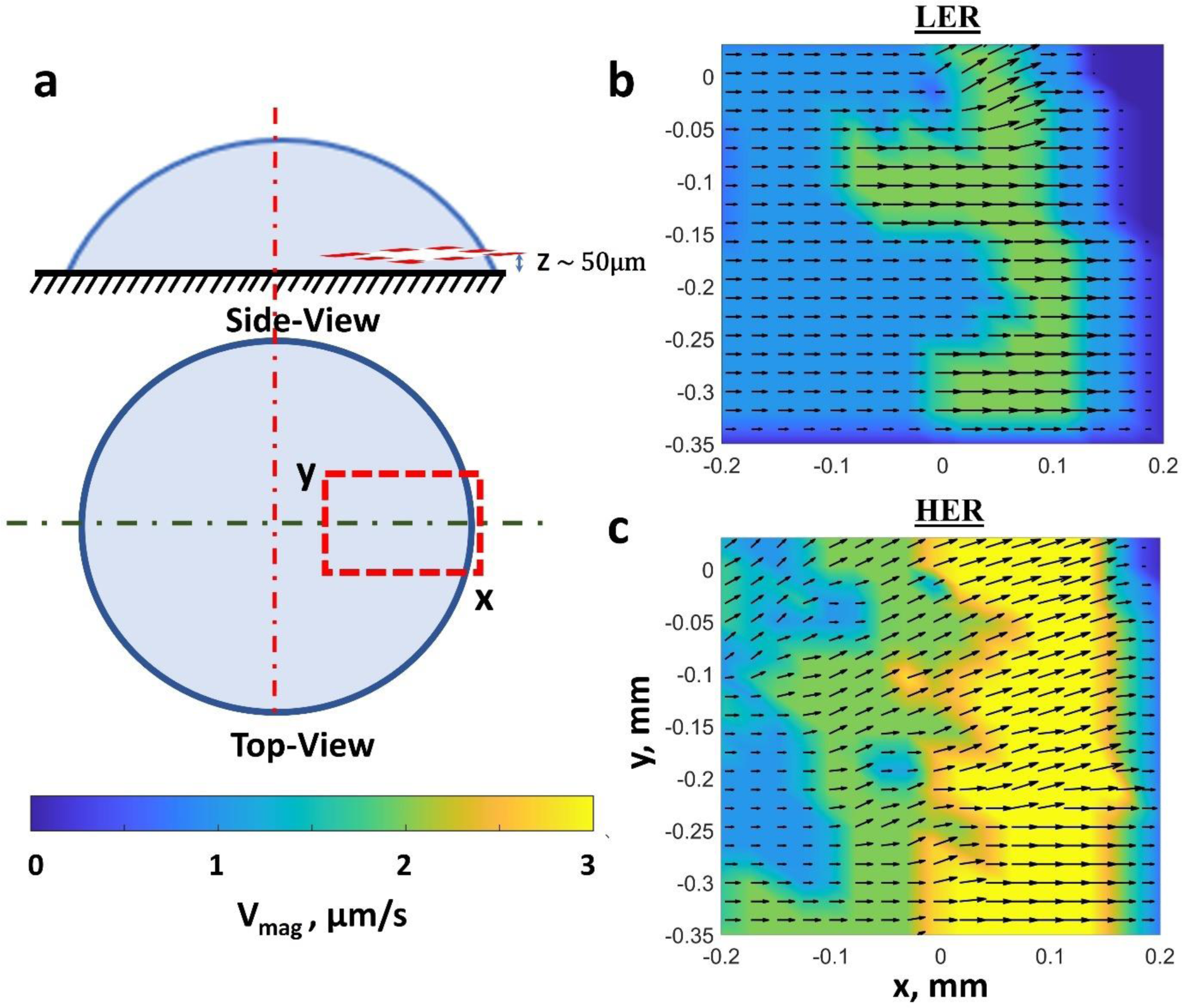
Micro-PIV measurements a) Schematic shows the plane of Micro-PIV measurements in side view and top view, and the scalebar for the velocity contour shown in b & c. b) shows the time average velocity vectors for LER case and c) HER case. The measurements are taken at a plane near the edge at 50 μm from the substrate surface at t/t_f_∼0.1.

The time-averaged velocity vectors shown in Fig. 4b&c represent the flow field for 5 seconds at t/t_f_∼0.1. The velocity contour represents the magnitude of time-averaged velocity at the given plane. For the HER case, the maximum velocity is around 3 μm/s, whereas for the LER case, the maximum velocity is approximately 2 μm/s. The height averaged velocity for the droplet drying in CCR mode can be given by, 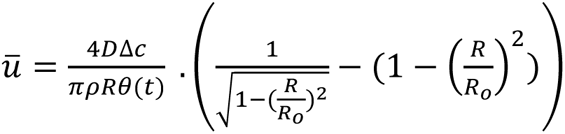 [63]. θ(t) is the instantaneous contact angle, and R is the radius at which velocity is measured. Near the contact line, the velocity can be approximated as 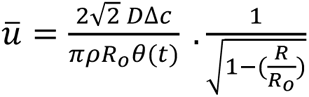 [63]. The velocity values obtained from these expressions align in order of magnitude with those measured experimentally. The magnitudes of outward velocity are greater in HER due to the higher mass flux than in LER. The maximum velocity occurs near the contact line for a pure solvent droplet, evaporating in pinned mode. In the SRF droplets, the location of maximum velocity is shifted inside due to edge deposits. The internal flow dynamics strongly influence the deposition of bacteria and other solutes. The local variation in solutal deposition and NaCl concentration alters the precipitation dynamics.

### 3.4. Precipitation dynamics

As the droplet undergoes evaporation, the contact angle steadily decreases, transitioning the shape from a spherical cap to a flat, thin film. Concurrently, water vapour escaping from the droplet interface increases solutal concentration over time, elevating salt, mucin, and bacteria concentration levels within the droplet. Eventually, the salt concentration reaches a saturation level. To initiate precipitation, the dissolved Na^+^ and Cl^-^ electrolyte concentrations must achieve a state of supersaturation, marking the onset of stable crystal nucleation and growth. In the case of a levitated droplet, NaCl crystal nucleation typically occurs at a supersaturation level of 2.31[64]. However, for a droplet on a surface, crystal nucleation might occur at a lower supersaturation level, facilitated by other solutal deposits serving as heterogeneous nucleation sites. Figures. 5a&b illustrate crystal growth at various time points. The rate of NaCl concentration increase under HER conditions would surpass that in LER conditions as the evaporation rate is higher. Thus, in HER droplets, crystals are observed to nucleate simultaneously from multiple sites, with the initial nucleation majorly occurring at the inner edge of the coffee ring where NaCl concentration is higher. Nucleation from the inner region also occurs at later times. In the case of HER, crystallisation commences at t/t_f_ = 0.92. The predominant crystal shapes are equiaxed dendritic structures with branching. Crystal growth extends outward from the nucleation sites. The growth rate reduces when crystals grow in the opposite direction. Crystal growth ceases when two crystal growth fronts meet.

**Figure 5.**
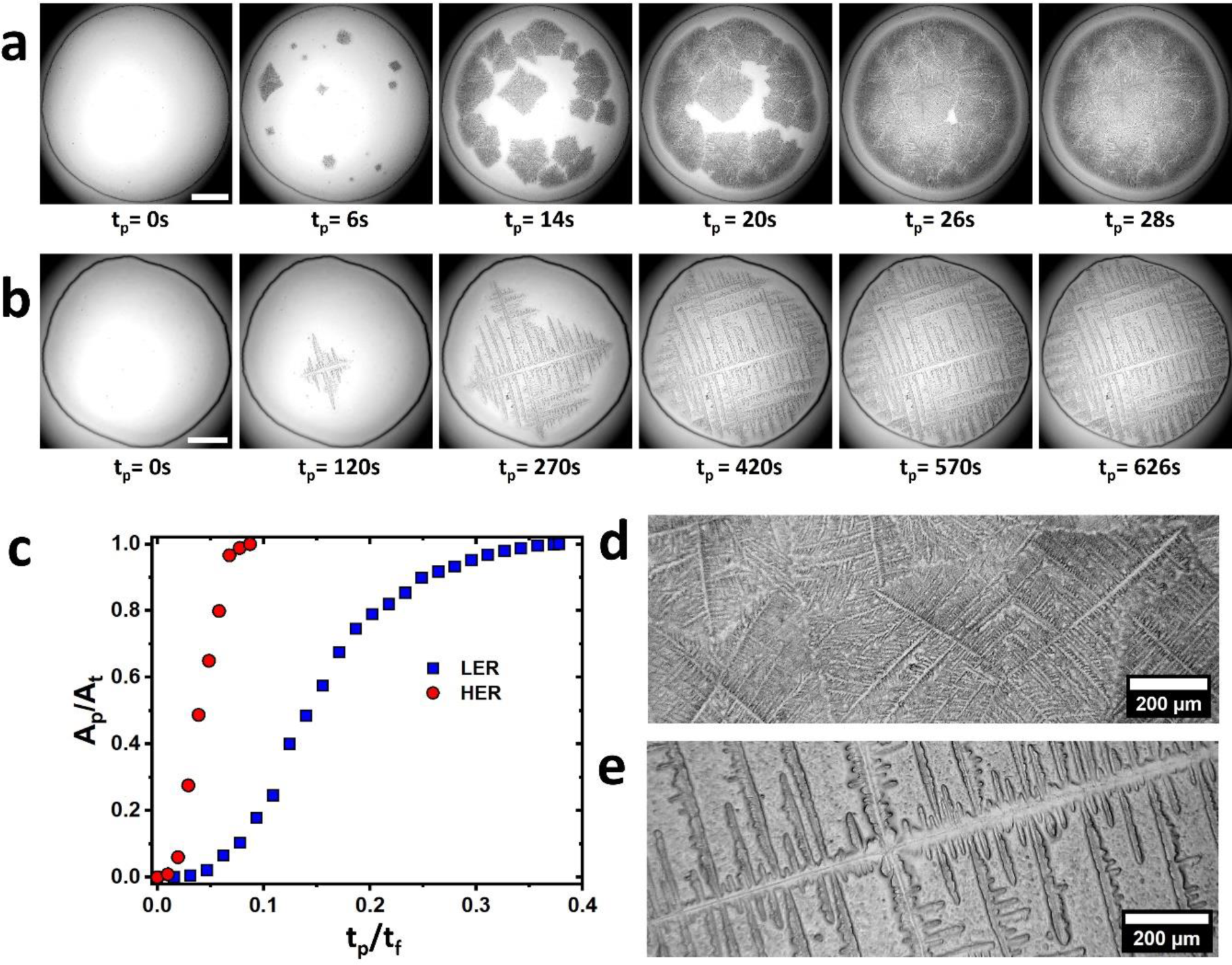
Precipitation dynamics: a) Droplet top view images at different time instances show the progress in precipitation for HER case, where t_p_=0 at the instant just before the first nucleation of the crystal becomes visible. The scale bar corresponds to 500 μm. b) Temporal top-view images of LER case during crystal growth c) Plot shows the ratio of instantaneous area precipitated, A_p_ to total area precipitated, A_t_ with normalised time t_p_/t_f_ for HER and LER cases. d) & e) shows the crystal structure of d) HER and e) LER deposits. The crystals in HER case deposits are multiple thin equiaxed dendritic structures with fine branches. In contrast, in the LER case, the crystals are much thicker than in the HER case and originate from a single equiaxed dendritic nucleate.

Under LER conditions, crystallisation occurs at t/t_f_ = 0.63. The reduced evaporation rate leads to a slower increase in NaCl concentration within the thin film. Across all repetitions, crystal growth consistently initiates from a single nucleation site located at the central region. This could be possible due to the water sorption by dense mucin deposits at the edge, resulting in lower net mass outflux than the inner regions. Thus, salt concentration first reaches supersaturation at the inner regions, contrary to HER, and crystal nucleates first in the inner region in the LER case. Na^+^ and Cl-ions diffuse towards the growing crystal as crystals grow. The depletion of NaCl in the growing crystal and the slow increase in NaCl concentration of LER prevents the formation of stable nucleation sites in other locations. The equiaxial dendritic crystals grow outward and branch. Notably, in the case of LER, the dendritic crystals and their branches are thicker than HER. Crystallisation takes approximately 700 seconds to complete under LER conditions, whereas it only takes around 28 seconds under HER conditions.

Figure 5c shows that in the case of HER, the crystallisation process occurs for a duration of 0.1t_f_, whereas it took around 0.4t_f_ in the case of LER. The rate of crystallisation, dAp/dt variation with non-dimensionalised time t_p_/t_p, t_ is shown in supplementary figure 2c, where t_p, t_ is the total time for crystallisation. The crystallisation rate increases from the onset of the first crystal nucleation and attains a maximum of around t_p_/t_p, t_ of 0.3 to 0.4 for both cases. The precipitation rate reduces after that due to dissolved Na^+^ and Cl^-^ depletion. The crystal’s growth rate increases with the increase in the rate of NaCl concentration change. The local concentration variation of NaCl depends on the net evaporative flux during crystallisation. Even though the initial mass flux in LER is of the same order as that of the HER case, near dryout mass flux could have drastically reduced, resulting in one order reduction in crystal growth rate in LER. During crystallisation, solutal Marangoni convection-induced flow occurs towards the crystal[65,66]. This flow could carry other solutes and bacteria along with the salt ions. Thus, the final bacterial distribution in the deposits could depend on the internal flow dynamics and precipitation dynamics.

### 3.5. Pathogen distribution

Bacterial deposition is initiated at the droplet edge since the evaporation-induced flow brings the bacteria to the contact line. The outward capillary flow propels bacteria and mucin towards the edge, where they eventually settle. Agrawal et al.[67] demonstrated that bacterial depositions in drying sessile droplets differ from those of homogeneous inert spherical polystyrene particles. The presence of mucin could introduce additional complexities to bacterial deposition. A detailed study is required in the case of LER to understand the effect of bacterial motility on final deposition.

This study captures fluorescently tagged PA using a confocal microscope to obtain the final pathogen distribution. Images at various z locations are superimposed to generate an overall pathogen distribution, as illustrated in Figures 3a and 3b for HER and LER cases. HER deposits exhibit dense pathogen concentrations at the coffee ring compared to LER, possibly due to higher capillary flow, as shown in Fig. 2. The edge deposition is more diffused and broader in the LER case. No significant variations are observed in the inner regions between the two cases. The distribution of pathogens along a radial direction from edge to edge is presented in Figure 3c. The mean gray value represents the average intensity of the fluorescence signal within a band of width 100 μm. The mean grey value peaks at the edge, indicating a higher pathogen density. The peak mean gray value at the edge is higher for HER than LER. The edge deposition extends for x/D ≈ 0.1 for HER and x/D ≈ 0.15 for LER. In the inner regions, the mean grey value of LER is slightly higher than that of HER. The distribution of pathogens within the deposit is a crucial factor influencing its viability, as desiccation levels can vary based on the deposit’s location[68,69].

**Figure 6.**
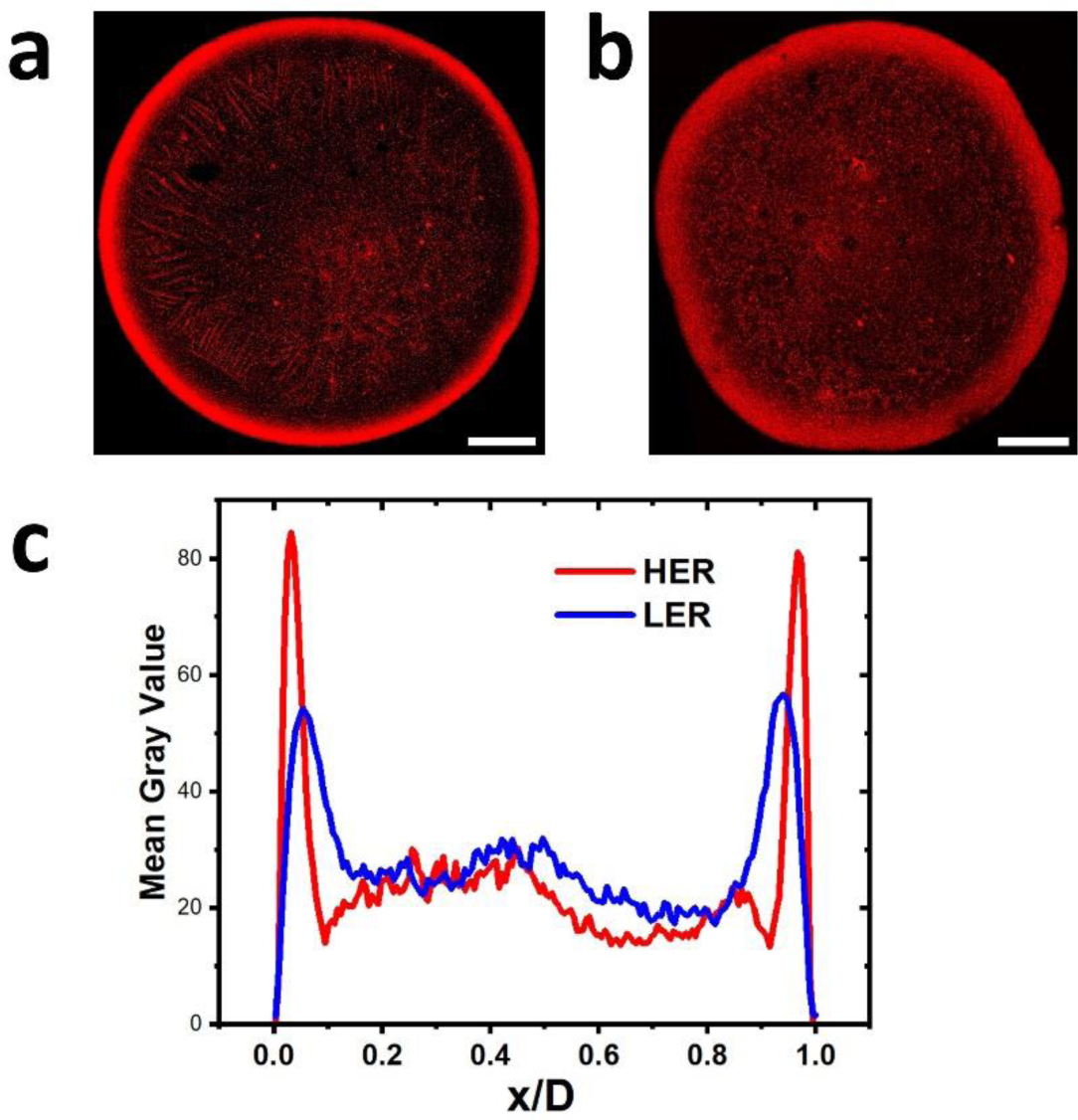
Pathogen distribution in dried deposits. a) superimposed z-stack confocal images show pathogen (red signal) deposition at HER conditions b) at LER conditions. Scale bar indicates 500μm c) Pathogen distribution along the deposit, shown by variation in mean gray value along the horizontal bar of width 100 μm through the centre of the deposit from the left edge to the right edge, where x=0 at the leftmost edge and x=D at the rightmost edge.

### 3.6. Bacterial Viability

**Figure 7.**
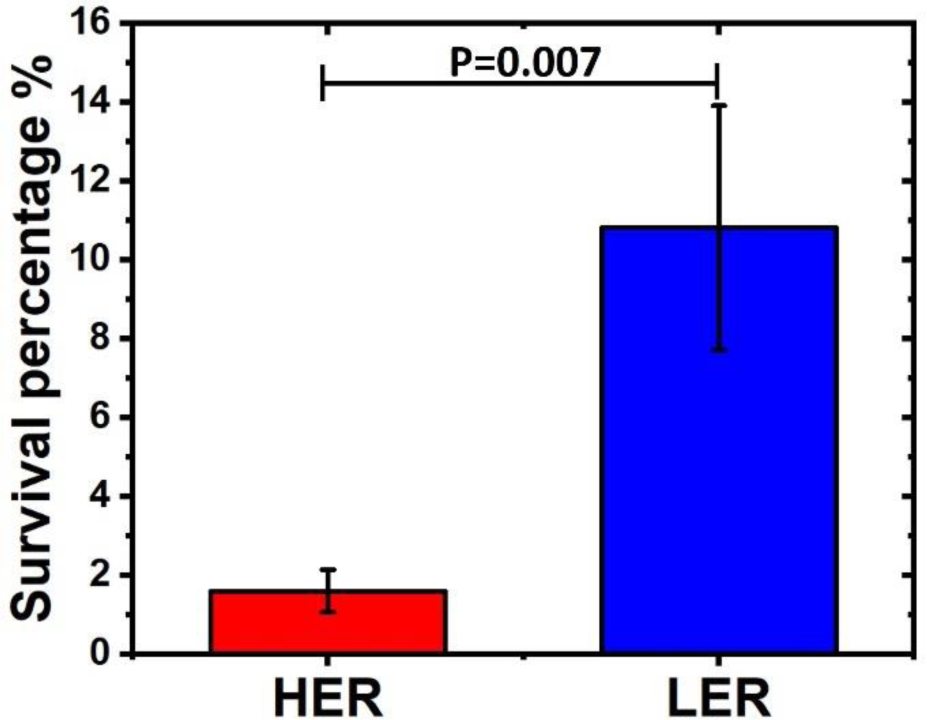
Viability of the *Pseudomonas Aeruginosa*

The culture-based bacterial viability studies consistently demonstrated that, for each repetition, PA survived at least five times more under LER conditions than HER conditions. The average survival percentage under high evaporation rate conditions was 1.6, but it increased significantly to 10.8 under low evaporation. This elevated survivability in LER conditions can be attributed to the lower desiccation stress, given the slower rate of water loss from bacteria. Here, the viability study shows that at least 85% of bacteria die under various stresses when exposed to ambient conditions for five hours. The bacteria buried within the deposits and shielded from the ambient environment would experience additional protection from desiccation compared to exposed bacteria. Despite the fact that denser pathogen distribution results in high viability[68,69] LER case has shown higher viability than HER. We strongly suspect that increased water sorption and reduced desorption could occur at high relative humidity, as reported by Björklund et al.[42].

Consequently, the hydrated mucin could act as an additional protective barrier, further shielding the bacteria from desiccation stress under high RH conditions and contributing to higher survivability, as depicted in the figure. 8. Here, we hypothesise mucin hydration as a potential reason for elevated bacterial survivability on inanimate surfaces at LER conditions. The validity of the mucin hydration is verified at higher humidities of 80% RH and 90%RH. At 80% RH, the crystallisation has not started even after 24 hours. When the deposits that are crystallised at LER and HER conditions are exposed to 90% RH, the crystals and precipitates dissolve, indicating water sorption in the precipitates. The detailed dynamics at elevated RH can be studied in the future. Increasing salt concentration in evaporating droplets could also induce osmotic stress on bacteria. The response to the combined effects of osmotic and desiccation stress can vary among bacterial species. However, PA has been observed to undergo an osmo-adaptive process, allowing it to survive in hyperosmotic stress conditions[70,71]. Additionally, bacteria in sessile droplets may encounter shear stress due to internal flow. Here, shear stress on PA is much less at very low velocities of a few micrometres per second. Therefore, in the current study, desiccation stress is suspected to be the significant factor determining PA survival.

**Figure. 8:**
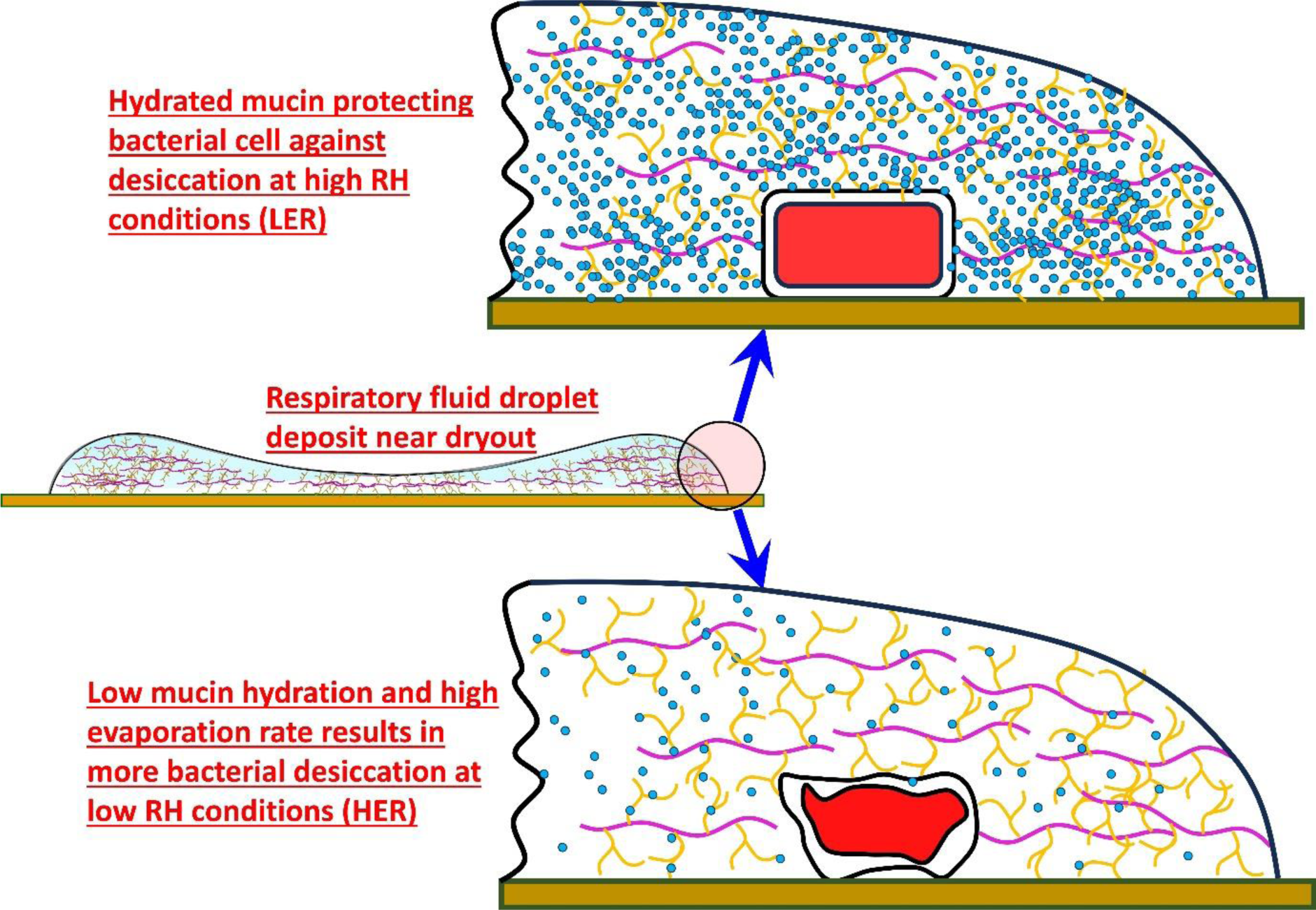
The schematic illustrates how well-hydrated mucin deposits could shield bacteria from desiccation in the case of LER, whereas the bacteria in the case of HER undergo higher desiccation in the presence of less-hydrated mucin deposits.

## 4. Conclusion

The PA-laden SRF droplet on the glass substrate exhibited faster evaporation at low RH conditions and slow evaporation at high RH conditions. A significant variation in internal flow velocities is observed between the LER and HER cases. The increment in solutal concentration reduced the evaporation rate towards the end of evaporation, particularly in the case of LER. The onset of crystal nucleation is much delayed in the case of LER compared to HER. The drastic delay in crystal nucleation could be attributed to the water-sorption characteristics of mucin. Crystal nucleation occurs at random locations near the droplet edge in the case of HER.

In contrast, crystals always nucleate from the central regions in the case of LER due to the differences in net interfacial mass flux created by the hydration of thicker mucin deposits near the droplet periphery. The PA distribution between the two cases shows differences at the periphery. HER showed denser peripheral deposition, while peripheral deposits are comparatively diffused in LER. On average, more than 10% of PA survived in the case of LER, whereas only less than 2% of PA survived in the case of HER. Previous studies showed that denser pathogen distribution results in more survival[68,69]. Even though pathogens are packed denser near the droplet edge for HER than LER, the viability of LER is higher than HER. The hydration of mucin due to water sorption is hypothesised as a primary reason for differences in evaporation, efflorescence, and survivability. Even though water sorption is not directly measured in this study, the literature and the current results strongly imply the presence of water sorption. A scope for many further investigations arises from this study, primarily on the water sorption and desorption kinetics in respiratory fluid deposits. The study’s implications can also apply to similar pathogens on biofluid droplets containing hygroscopic constituents like mucin. The study emphasises the importance of sanitising inanimate surfaces in frequently used community facilities, such as hospitals, public transport, restaurants, and schools, especially in highly humid conditions.

## Note

The authors declare no competing financial interests.

## Credit Statement

Conceptualisation- SB, DC; Methodology- AR, KP, SB, DC; Validation- AR, KP, SJ; Formal analysis- AR, KP, SJ; Investigation- AR, KP, SJ; Resources-SB, DC; Data Curation- AR, KP, SJ; Writing Original Draft- AR; Writing Review & Editing- AR, KP, SJ, SB, DC; Visualization- AR, KP, SJ; Supervision- SB, DC; Project administration- SB, DC; Funding acquisition- SB, DC.

Abdur Rasheed-AR, Kirti Parmar-KP, Siddhant Jain-SJ, Dipshikha Chakravortty-DC, Saptarshi Basu-SB.

## Acknowledgements

The authors thank the central facility at the microbiology and cell biology department at IISc for access to confocal microscopy. SB: PW Chair Professorship, the support by Serb-SUPRA (Scientific and Useful Profound Research Advancement) through project no.

SERB/F/10572/2021-2022 is thankfully acknowledged, DC: DAE-SRC fellowship, ASTRA-Chair fellowship, TATA Innovation grant, DBT-IOE partnership grant.

## Data availability statement

All data are available in the main text or supplementary materials. All materials and additional data are available from the corresponding author upon request.

