## Supplementary data for "Weather-related changes in the dehydration of respiratory droplets on surfaces bolster bacterial endurance"

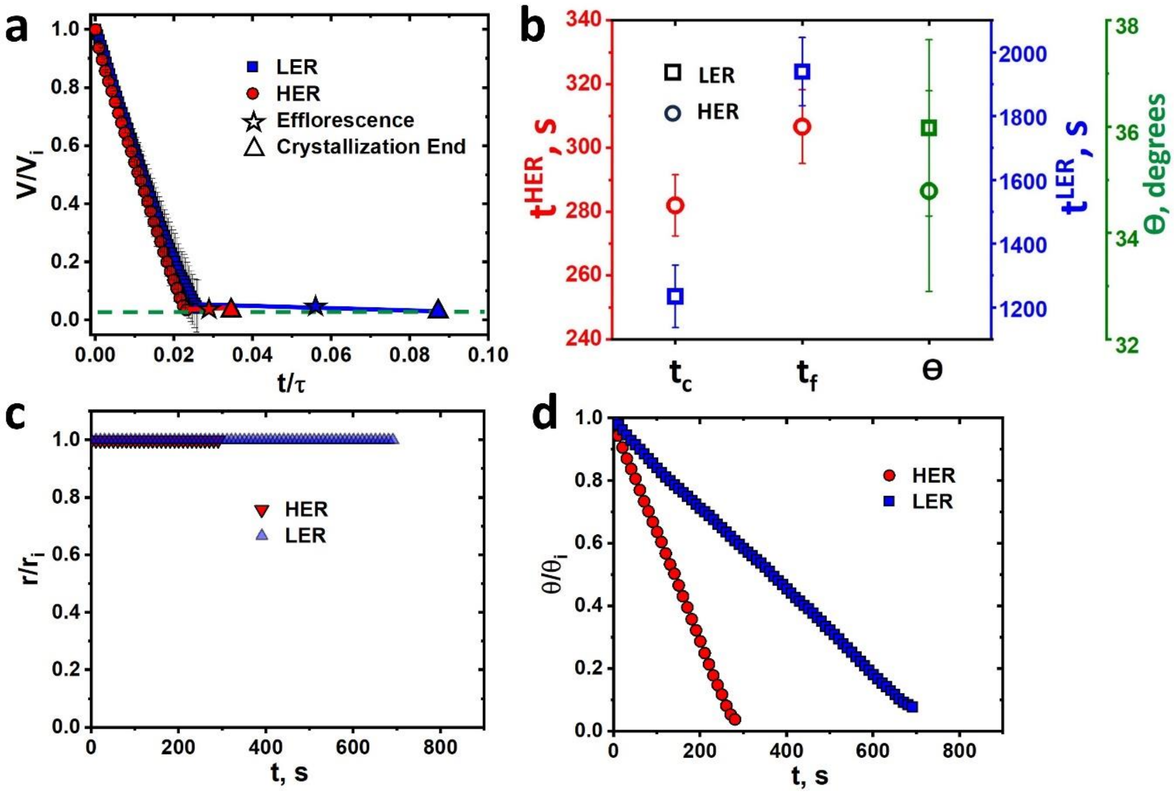


**Fig: S1.a) Normalized volume $V/V_i$ change with nondimensionalized time $t/\tau$ where $\tau$, is the characteristic time for diffusive evaporation. b) Plot shows the time taken for crystallization initiation $t_c$, time taken till the end of crystallization, $t_f$ and the initial contact angle c) Plot shows the normalized contact radius variation with time. d) Plot shows the normalized contact angle variation with time**

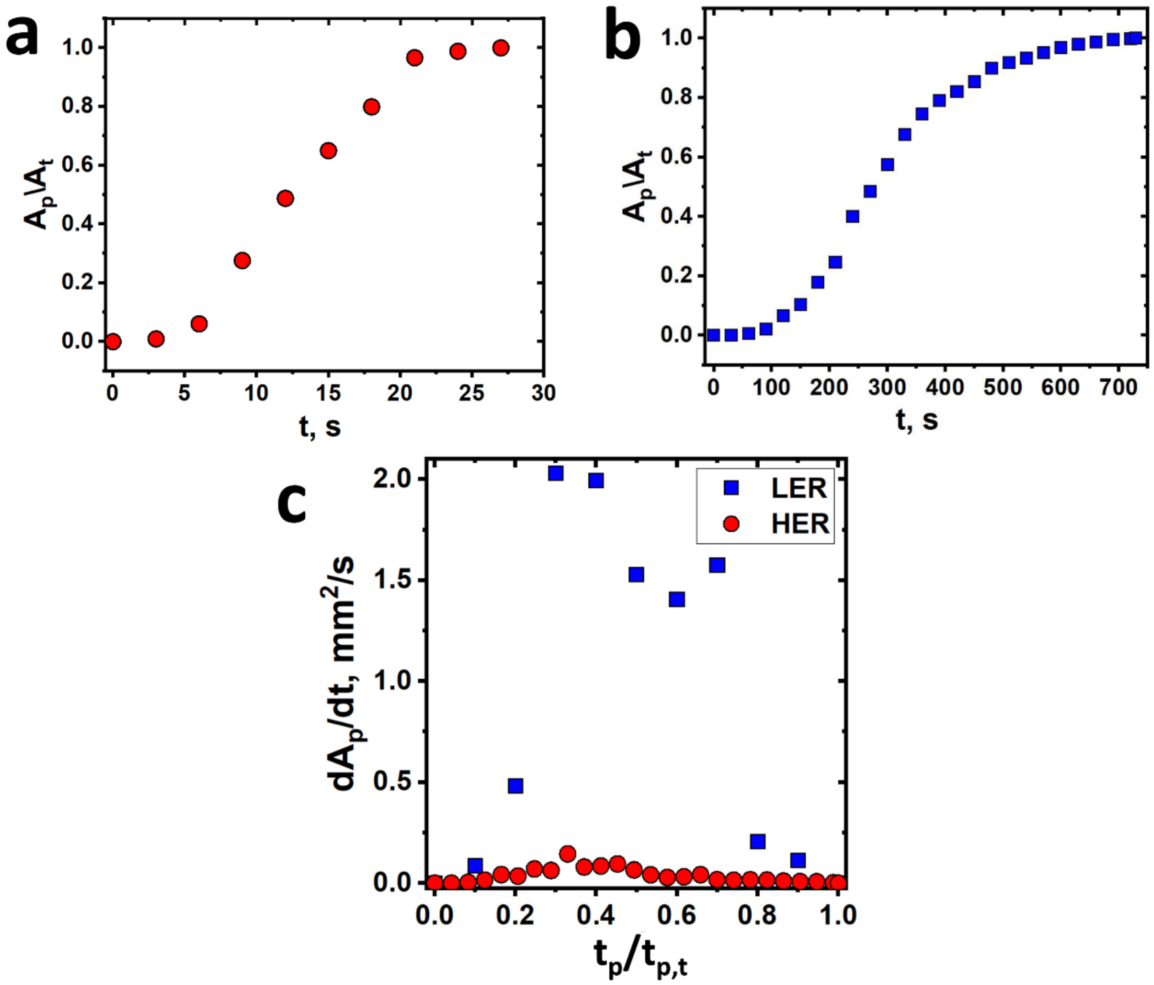


**Fig. S2. Normalized precipitation area variation with time for case a) HER and b) LER. C) Shows the precipitation rate $dA_p/dt$ variation in $mm^2/s$, with nondimensionalized time $t_p/t_{p,t}$, where $t_p$ is the time since efflorescence and $t_{p,t}$ is the total time for precipitation.**
